# Modelling Phenotypic Assay Noise for Multivariable Biomarker Discovery using Deep Neural Networks

**DOI:** 10.64898/2026.11.19.767940

**Authors:** James J. Bannon, Katy L. Scott, Farnoosh Khodakarami, Marc Delgado Sanchez, Robert F. Gruener, R. Stephanie Huang, Benjamin Haibe-Kains

**Affiliations:** Princess Margaret Cancer Centre, University Health Network, Toronto, Ontario, M5G 2C1, Canada; Medical Biophysics, University of Toronto, Toronto, Ontario, M5G 2C4, Canada; Structural Genomics Consortium, Toronto, Ontario, M5G 1L7, Canada; Department of Computer Science, University of Toronto, Toronto, Ontario, M5S 2E4, Canada; Vector Institute for Artificial Intelligence, Toronto, Ontario, M5G 0C6, Canada; Department of Experimental and Clinical Pharmacology, College of Pharmacy, University of Minnesota, Minneapolis, Minnesota; Masonic Cancer Center, University of Minnesota, Minneapolis, MN 55455, USA Department of Experimental and Clinical Pharmacology, College of Pharmacy, University of Minnesota, Minneapolis, Minnesota; Bioinformatics and Computational Biology, University of Minnesota, Minneapolis, Minnesota; Universitat de Barcelona, Biomedicine; KTH, Stockholm

**Keywords:** Deep Learning, Pharmacogenomics, Treatment Response, Noise Robustness, Personalized Medicine, Cancer

## Abstract

Accurately predicting cancer drug response remains a major challenge in precision oncology. Although some therapies are guided by clinically-approved biomarkers, these markers often yield limited predictive value and are unavailable for many drugs. Artificial Intelligence and Machine Learning approaches trained on large-scale cancer cell line pharmacogenomic screens offer a promising route to more comprehensive response prediction, yet they generally assume that phenotypic readouts are measured without error. This assumption is problematic because viability-based drug response assays are inherently noisy, potentially causing models to learn experimental artefacts rather than biologically meaningful determinants of sensitivity. Here, we show that phenotypic measurement noise is present in four widely used cancer cell line screening datasets and introduce Deep Concordance Index Networks (DeepCINET), a deep learning framework that explicitly accounts for this noise. Across datasets, incorporating this noise-aware ranking strategy improves generalization across experimental conditions and to previously unseen samples. DeepCINET outperforms baseline models in multiple settings, with particularly notable gains in prioritizing the most drug-sensitive cell lines. Together, these results establish phenotypic assay noise as a critical and previously overlooked source of error in drug response modelling and provide a principled framework for noise-aware multivariable biomarker discovery.

**Key Points:**

1. We show that measurement noise exists in phenotypic measurement in common cell line viability screens and that accounting for this noise increases across-screen consistency.
2. We introduce a hyperparameter δ that defines pairs of cell lines as valid pairs (resp. invalid) if their AAC measurements to a given drug differ by more than (resp. less than) a value δ.
3. We develop a novel deep learning architecture called DeepCINet to predict drug response from data that learns to order cell lines by looking only at valid pairs in the training set.
4. DeepCINet shows improved predictive performance in cross-screen analysis on both seen cell lines as well as new unseen cell lines.

## Introduction

Predicting which patients will respond to a given cancer therapy remains an open problem of profound clinical importance. For some therapies there exist *biomarkers*, collections of (molecular) measurements that have been associated with a given treatment outcome and which can be used to guide treatment decisions[1]. For example, the presence of an EGFR mutation in non-small cell lung cancer is often predictive of response to tyrosine kinase inhibitors (TKIs), and is widely used as a clinically-approved biomarkers for these targeted therapies[2]. While the discovery of predictive biomarkers has greatly improved the precision with which treatments are selected, the predictive value of existing biomarkers is often limited. For instance, patients with an EGFR mutation do not uniformly benefit from the TKI erlotinib [3]. Additionally, not every drug has a biomarker that can be used to determine the potential for patient response [4].

Artificial intelligence (AI) and machine learning (ML) methods are increasingly being used to predict cancer drug response [5,6]. Conceptually, these models function as multivariable biomarkers that capture complex tumour biology more comprehensively than single molecular markers. Most biomarkers are developed using large *in vitro* cancer cell line (CCL) viability screens, where drug-dose response curves are summarized as IC50 or area above the dose-response curve (AAC) [7]. However, we and others have shown that these phenotypic readouts are inherently noisy [7,8]. This variability has been largely overlooked in prior AI/ML studies, despite the risk that models may learn experimental artefacts rather than biologically meaningful response patterns. In particular, when the measured responses of two cell lines differ only within the expected range of assay noise, their relative ordering is uncertain and may not reflect a true biological difference in sensitivity. We therefore hypothesized that explicitly accounting for assay noise during model training would improve the identification of genuine genotype-phenotype relationships.

Here we present Deep Concordance Index NETworks (DeepCINET), an approach to explicitly account for phenotypic measurement noise during training of drug response prediction models. DeepCINET is a unique combination of two complementary techniques from the learning-to-rank literature: ranking using pairwise comparisons made with siamese neural networks as presented in RankNet[9] and ignoring potentially noisy comparisons[10]. To our knowledge, DeepCINET is the first application of those techniques to drug response prediction. We first show that this noise is present in four cancer cell line viability screens widely used to train and evaluate AI/ML models. We then demonstrate that a noise-aware version of DeepCINET frequently outperforms an otherwise identical version that does not account for measurement noise. Finally, we show that DeepCINET can outperform machine learning baseline models when evaluated on new samples. Notably, these gains are especially pronounced for identifying the most drug-sensitive cell lines, a task of particular practical relevance for biomarker discovery and treatment prioritization.

## Results

We developed DeepCINET, a novel model for drug response prediction that accounts for phenotypic noise. Specifically, DeepCINET is implemented as a siamese neural network architecture [11] that processes pairs of cell lines simultaneously. For each pair, the molecular omics profiles of the two cell lines are passed through a shared neural network to produce sensitivity scores. Following the RankNet framework [9], the model is trained on pairs of cell lines and uses a binary cross-entropy loss to encourage the model to assign a higher score to the cell line exhibiting greater sensitivity to the selected drug. A key distinction between DeepCINET and RankNet is the introduction of a non-negative *separation parameter* δ, adapted from Ohue et al. [10]. Two cell lines are considered to form a valid training pair if and only if the absolute difference between their measured viability values exceeds experimental noise, represented by δ. Consequently, DeepCINET is trained exclusively on these valid pairs, thereby reducing the influence of potentially noisy or biologically ambiguous comparisons (Figure 1a, Figure 2d).

**Figure 1:**
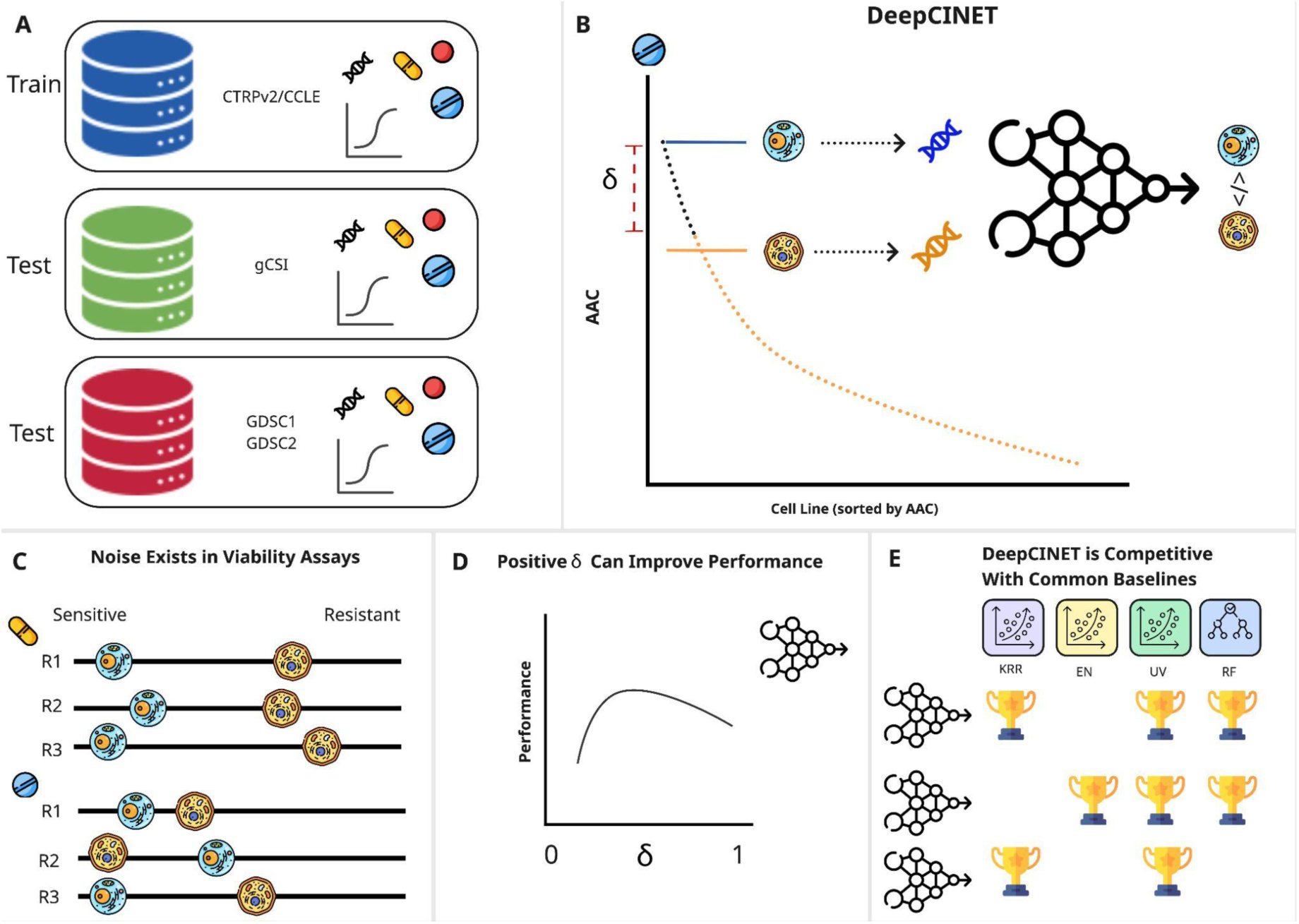
DeepCINET Architecture and Study Overview. (A) The dataset used. We used four common cancer cell line screens for our work: CTRPv2, gCSI, GDSC1, and GDSC2. All four have dose-response curves and omics measurements for hundreds of cell lines and a wide range of drugs. (B) The DeepCINET approach. For a given drug, DeepCINET partitions a set of cell lines into *valid pairs*, which are pairs of cell lines with AAC measurements that differ by more than a fixed hyperparameter δ. DeepCINET is then trained to use omics measurements from those cell lines to rank only the valid pairs in proper order. The indicated cell lines form a valid pair. For the top cell line (blue) any cell line with AAC with an acc below the delta mark (yellow dots) would form a valid pair with the blue cell line. Panels (C-D) show the study outline. (C) Demonstrates the potential pitfall of ignoring phenotypic measurement noise; for some drugs, which cell line is more sensitive to the given drug may change across replicates. (D) We show that setting the hyperparameter δ to a strictly positive value improves the performance of the model. (E) Finally, we show that DeepCINET is competitive with strong baselines and in some cases can radically improve generalization performance.

**Figure 2:**
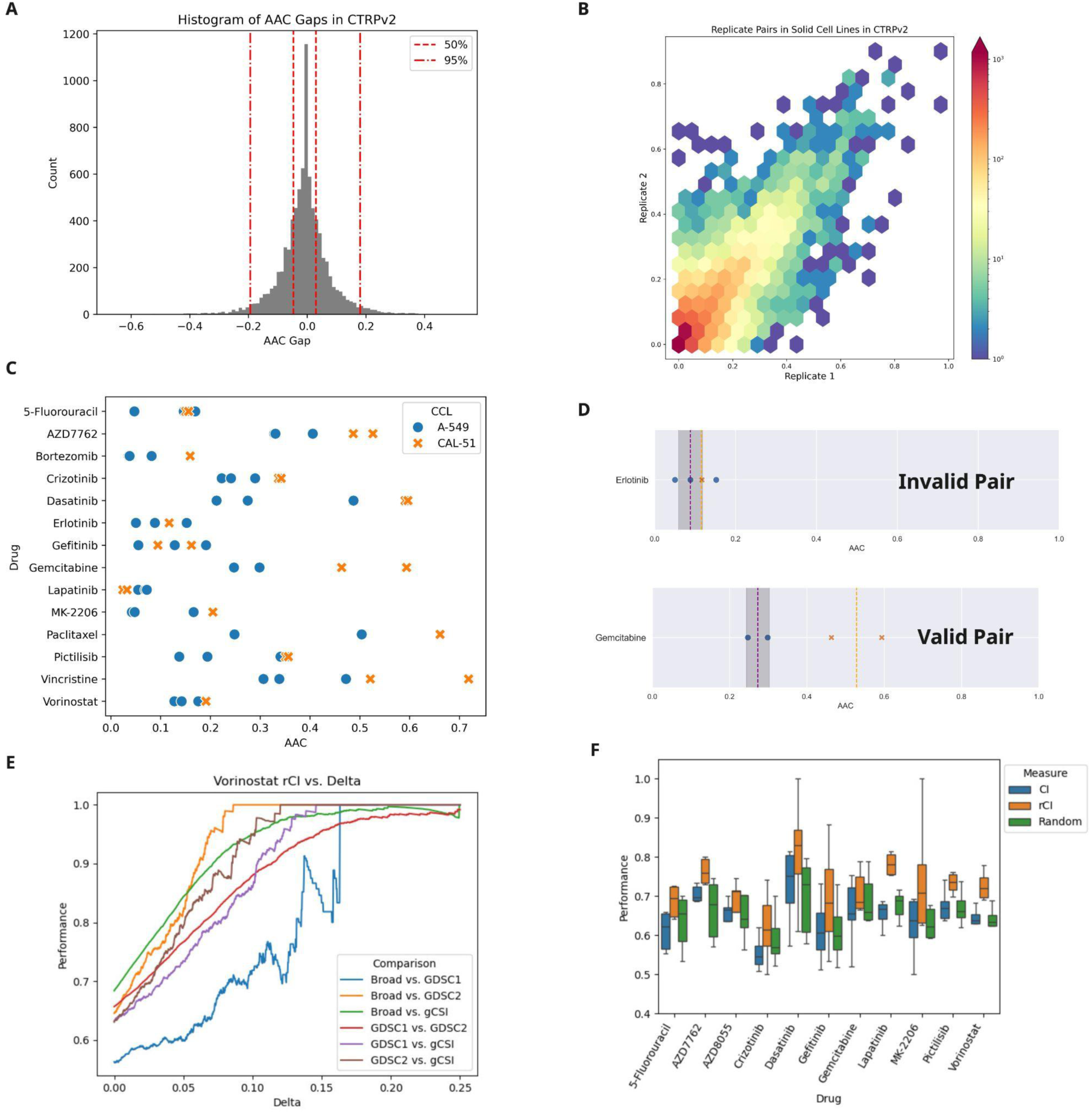
Phenotypic noise exists in the assay. We demonstrate that phenotypic measurements of drug response are noisy by displaying the distribution of differences in AAC for pairs of replicates in the CTRPv2 screen. **(A)** A histogram of the difference in AAC measurements for replicates in solid tissue cell lines. **(B)** A hexplot showing the density of replicate pairs in the CTRPv2 by the *absolute* differences in their AAC values. **(C)** Replicate AAC values for the drugs considered in two cell lines A-549 (blue dots) and CAL-51 (orange crosses). **(D)** An example of how the same two cell lines can form or not form a valid pair depending on the drug considered. The vertical bar is at the median AAC value for A-549 and the grey band spans a total area of the average absolute difference in CTRPv2 (i.e. δ ≈ 0. 05). The purple vertical line is at the median AAC for A-549 and the orange vertical line is the median AAC for CAL-51. For Erlotinib these cell lines would not be considered a valid pair as the median AAC values are not separated by more than δ. The same cell lines would be a valid pair for Gemcitabine. **(E)** Increasing the value of δ can increase the robust concordance index (rCI) in AAC measurements between screens. We show how there is a landscape in potential improvement as a function of δ. **(F)** We show that across all drugs increasing delta can improve drug response measurement consistency and that the removal of random pairs has no impact on performance.. We set =0.03 in all drugs for these comparisons.

Our study uses four cell line viability screens that are commonly used in the machine learning literature: the Cancer Therapeutic Response Portal v2 (CTRPv2) [12], the Genentech Cell Line Screening Initiative (gCSI) [13], and both releases of the Genomics of Drug Sensitivity in Cancer screen (GDSC1 and GDSC2 using the Syto60 and CellTiter-Glo assays, respectively) [14]. We augmented the viability measurements in CTRPv2 with the molecular measurements from the Cancer Cell Line Encyclopedia (CCLE) [15] to create our training set. We evaluated all trained models on the data from GDSC1, GDSC2 and gCSI in two scenarios: the performance on cell lines encountered during training - “seen” cell lines - and cell lines new to the model - “unseen” cell lines. In all cases we used RNA-seq data as the input to the model. Since there is some overlap in the RNA-seq data between GDSC1 and GDSC2, the independent evaluation on the test set is about the *phenotypic assay noise only*. We only trained and evaluated DeepCINet on solid tissue tumour cell lines.

### Quantifying the Noise in the Drug Response Data

To quantify the phenotypic noise in our four datasets we analyzed the distribution of differences in area above the dose-response curve (AAC) [14] measurements across replicates within each study. We found that the distributions are concentrated around zero with an average absolute difference between replicates around 0.05 in GDSC2, gCSI, and CTRPv2 and closer to 0.07 in GDSC1 (Table 1). These distributions also have heavy tails meaning that for the same drug-cell line pair the measured sensitivity values can be quite different (Table 1, Figure 2a 2b, Supplementary Figure 1). In fact, we see cases where the same drug-cell line combination can have an AAC measurements, which lie between 0 and 1, that differ by over 0.5 between replicates (Figure 2b, Supplementary Figure 1). This means that in one replicate a cell line may be measured as sensitive to a drug and in another it may be measured as resistant[16] (Figure 1c).

In addition to confirming the noise intrinsic to viability screens, our results corroborate our previous work on inconsistency between studies [7,8]. For example, the correlation between AAC measurements in the training set and the test sets is modest [17], with the value lying below 0.6 in most cases (Figure 3A-C left side, vertical bars). These inconsistencies affirm the need for AI/ML models to be tested on multiple independent datasets and highlight the need for caution in developing biomarkers from a single cell line screen.

**Figure 3.**
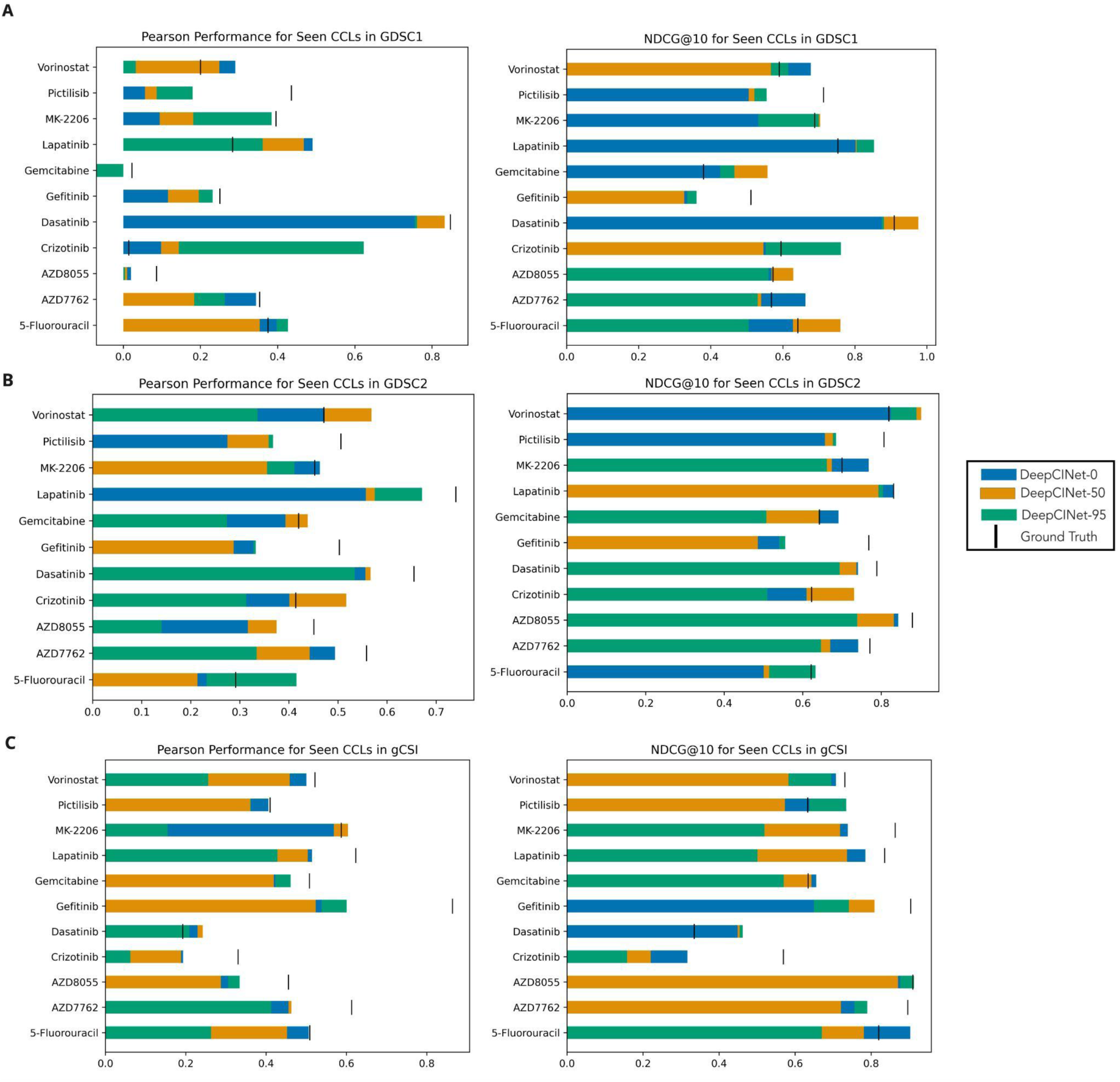
Impact of the Separation Parameter on Performance. We display the performance of DeepCINET with δ set to 0 (blue bars), and ◻ set to the 50th (orange) and 95th percentile (green) of the distribution of absolute differences between replicates. As pseudo ground-truth we use the measured AAC in the training set. The ground truth performance is marked as a black vertical bar. (A) The performance in terms of Pearson correlation and NDCG@10 in GDSC1. (B) The performance in terms of Pearson correlation and NDCG@10 in GDSC2. (C) The performance in terms of Pearson correlation and NDCG@10 in gCSI.

Noise in the phenotypic measurement across poses an issue for machine learning models as it can lead to spurious conclusions about the relative sensitivities between cell lines. This problem persists even if one takes summary measurement across replicates such as the mean or median. The summarized measurements for two cell lines can be within the range of typical replicate noise for the assay (Figure 2C, 2D). In cases such as this the conclusion that one cell line is more sensitive to a drug than another is dubious. It is not clear if the difference sensitivity measurements are due to genuine biological differences or artefacts of the assay and protocol being used. By considering pairs of cell lines with a non-trivial separation (i.e. δ>0) DeepCINET should avoid this potential pitfall.

### The Separation Parameter Improves Cross-Study Consistency

To determine if the separation parameter δ can in fact improve consistency between screens, we computed the robust concordance index (rCI) [18] between the AAC measurements for each drug in all pairs of screens (Figure 2F). Briefly, the rCI computes the number of pairs of cell lines that have AAC measurements that differ by at least δ in both screens and are ordered in the same way in both. Details on computing the rCI are provided in the methods section.

For a value of δ = 0. 03, we found that the rCI in every drug considered was higher in every drug (Figure 2F). The average concordance index (equivalent to rCI with δ = 0) is 0.65 in across all 11 drugs considered while average rCI δ = 0. 05 is 0.72. This corroborates that consistency across studies can be increased by ignoring potentially spurious pairs. We also investigated the dependence of the rCI value on δ by computing the rCI for all drugs and for all pairs of screens with δ ranging in 1,000 equally spaced values between 0 and 0.25 (Figure 2E, Supplementary Figure 4). While most drugs show the a pattern similar to Vorinostat (Figure 2E), namely, an increase in rCI as δ increases, this pattern is not universal. There are some drugs where δ does not impact the consistency across studies such as Gemcitabine when computing the rCI between CTRPv2 and gCSI. Also the linearity and monotonicity of the relationship appears to depend on both the drug and the pair of screens compared. Importantly the increase in rCI is due to the removal of noisy pairs. The removal of random pairs either degrades performance or leaves it unchanged (Figure 2F).

### The Separation Parameter Effectively Ignores Assay Noise

To assess whether the separation parameter δ can in fact help the model avoid fitting to assay noise, we trained a DeepCINET in three configurations: one with δ=0, one with δ set to the 50th percentile and one with 95th percentile of the absolute difference distributions in CTRPv2 (δ = 0. 0801 and δ = 0. 1878, respectively). We evaluated the performance of these models on the cell lines common to both the training and evaluation screen, i.e., cell lines already seen by the model during training. By focusing on seen cell lines we can directly evaluate our hypothesis that the inclusion of the positive delta parameter allows the model to learn genuine associations between omics measurements and phenotype.

We evaluated model performance using the Pearson correlation coefficient, as well as the Normalized Discounted Cumulative Gain at ranks 5 and 10 (NDCG@5 and NDCG@10). As a reference benchmark, we compared each DeepCINET configuration against a “ground truth” estimator derived directly from the actual AAC measurements. Specifically, this estimator uses the observed AAC value for each drug–cell line pair in the training dataset as the predicted response for the corresponding pair in the test dataset, providing an empirical “upper bound” for the predictive value of the models.

In every evaluation we see cases of improvement from including a positive delta. In GDSC1 we see improvements in Pearson correlation in 54% of cases (6 out of 11 drugs), and 73% (8 out of 11) and 82% (9 out of 11) in terms of NDCG@5 and NDCG@10, respectively. For GDSC2, a model with δ greater than zero led to an improvement in 82% of drugs (9 out of 11) when measuring performance by Pearson correlation, 45% (5 out of 11) of drugs when measuring performance by NDCG@10 and 64% (7 out of 11) of drugs in terms of NDCG@5. Lastly for gCSI the corresponding numbers are 54% (6 out of 11), 45% (5 out of 11), and 54% (6 out of 11) (Figure 3, Supplementary Figure 2).

### DeepCINET Can Generalize Better than Machine Learning Baselines

To assess how well DeepCINET generalizes to new samples we computed the performance of DeepCINET on the *unseen* cell lines in GDSC1, GDSC2, and gCSI, and compared it to four established machine learning models (ML), including kernelized ridge regression[19] (KRR), Elastic Net[20] (EN), Random Forests (RF)[21], as well as a univariable model that serves as a simulated single variable biomarker.

We introduce a fourth configuration for DeepCINET (referred to as DeepCINET-CV) where the separation parameter δ is tuned during training. For all models, including DeepCINET-CV, we chose hyperparameters using 10-fold cross validation. Choosing the separation parameter this way is relevant because the distribution of replicate values may not always be available. It is also important to evaluate how DeepCINET performs when δ is selected empirically from the training data as this is common practice in machine learning research. Cross validation selected a strictly positive value for δ in every drug considered except for AZD7762 (Supplementary Table 1).

We compared the best of the four DeepCINET configurations (DeepCINET-CV, DeepCINET-0, DeepCINET-50, and DeepCINET-95) to the best of the four ML baseliness. When measuring model performance using Pearson correlation we found that the best DeepCINET configuration outperformed the best baselines in 45% (5 out of 11), 64% (7 out of 11) and 45% (5 out of 11) of drugs in GDSC1, GDSC2, and gCSI, respectively. (Figure 4). The performance of DeepCINET relative to baselines improves when considering the ranking metrics. When measured in terms of NDCG@10 the best DeepCINET configuration outperformed the best baseline in 90% of the drugs (10 out of 11) in GDSC1, 82% (9 out of 11) in GDSC2, and 64% (7 out of 11) in gCSI. For NDCG@5 the values are higher for DeepCINET 73% (8 out of 11), 64% (7 out of 11) and 54% (6 of 11) in GDSC1, GDSC2, and gCSI, respectively (Figure 4, Supplementary Figure 3).

**Figure 4.**
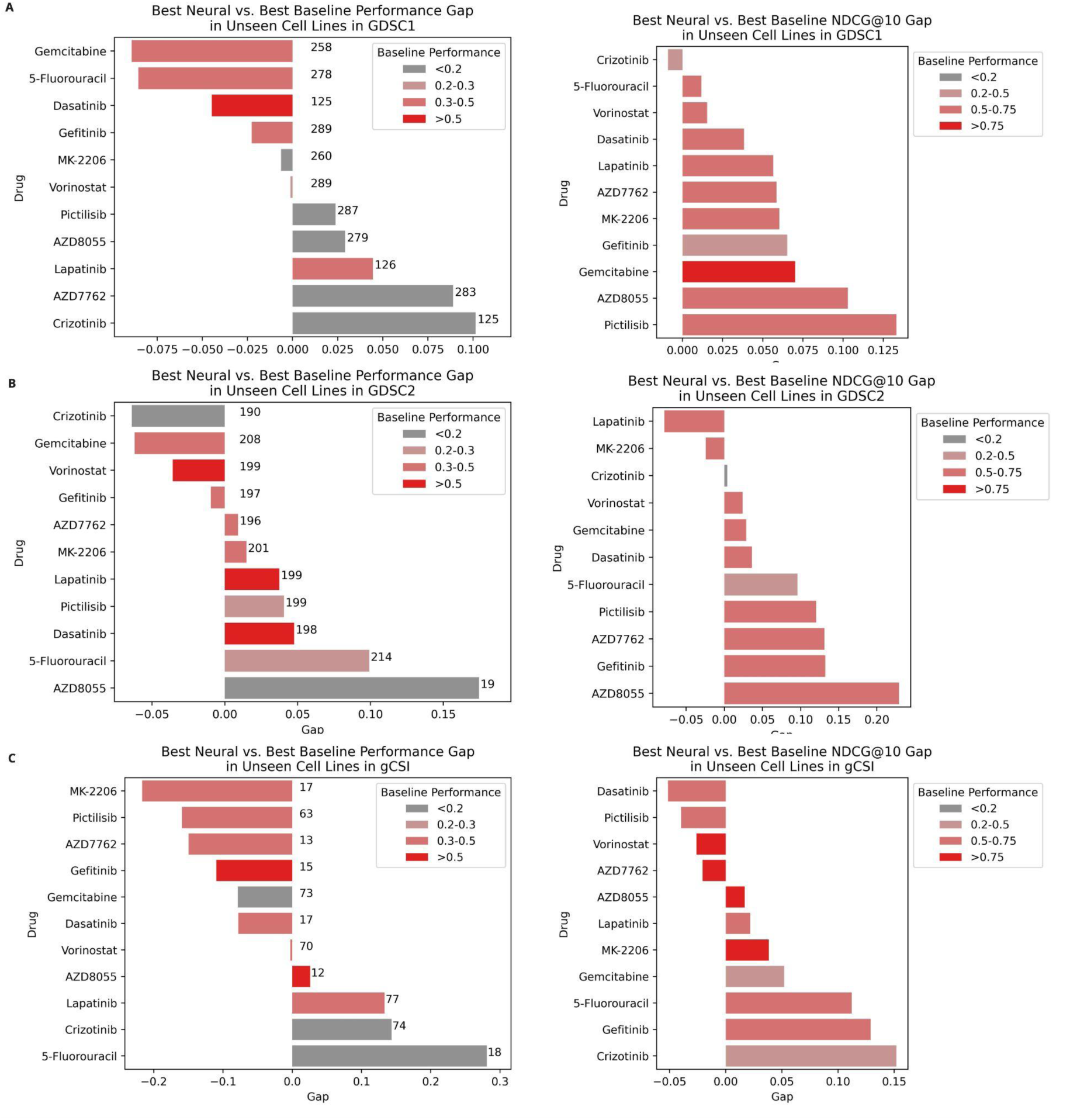
Comparison of Best DeepCINET vs. Best Baseline. Bar plots showing the performance gap (i.e., difference) between the Pearson correlation and NDCG@10 of the best neural model and the best baseline model in unseen cell lines in (A) GDSC1, (B) GDSC2, and (C) gCSI separately. The bars are colored by the performance of the best baseline model and bars in the Pearson plots are labeled by the number of samples used to evaluate model performance.

Interestingly, there are cases where DeepCINET is worse than the baselines in terms of Pearson correlation but performs better in terms of NDCG. These results suggest that while the model may not be predicting scores accurately across the entire test set, it is doing well at separating the cell lines with higher AAC values from those with lower AAC values.

For gCSI the number of unseen cell lines is relatively small (fewer than 20 samples) for many of the drugs considered. As such we focus on the results in unseen cell lines in GDSC1 and GDSC2. There are several drugs - AZD8055, Gefitinib, AZD7762, Pictilisib, Dasatanib, Gemcitabine, and Vorinostat - that show increased ranking performance for DeepCINet in both GDSC1 and GDSC2. To understand what distinguishes these drugs, we classified each of the considered drugs as ‘no effect’, ‘‘broad effect’ or ‘narrow effect’ as we have done in prior work [7]. Specifically we classified drugs using the distribution of their AAC values in solid cell lines in CTRPv2. A drug was categorized as having no effect if there were fewer than 5 cell lines with an AAC over 0.2, ‘narrow’ effect if the mean absolute deviation of their AAC values was below 0.13, and ‘broad effect’ otherwise. Of the drugs that show ranking improvements in both GDSC1 and GDSC2, all are classified as ‘narrow effect’ with the exceptions of Dasatanib and Gemcitabine (Supplementary Table 2). Many of these drugs have known targets; AZD8055 is known to be an mTOR inhibitor [22], Gefitinb is [23] an EGFR inhibitor, AZD7762 is a [24] checkpoint kinase inhibitor, Pictilisib is a PI3K inhibitor [25], Dasatanib is a tyrosine kinase inhibitor (TKI) for the SRC gene [26,27], and Vorinostat is an HDAC inhibitor [28].

Finally, in cases where the best DeepCINET configuration is worse than the best baseline, the difference is small in GDSC1 and GDSC2. In terms of Pearson correlation, the best baseline outperforms the best DeepCINET in Gemcitabine in GDSC1 by approximately 0.075 and by approximately 0.05 in Crizotinib in GDSC2. Thus, the separation parameter allows the DeepCINET models to remain competitive with machine learning baselines in terms of Pearson correlation while also giving improvements in terms of ranking metrics.

## Discussion

We have presented DeepCINET, a novel approach to training deep neural networks for drug response prediction. The core innovation of DeepCINET is the combination of techniques from the learning to rank literature [9,10] and their application to drug response prediction. To justify the need for DeepCINET, we have expanded on our previous work [7,8] and have shown that there is indeed substantial phenotypic measurement noise in four cell line viability screens commonly used in the machine learning literature. We then showed that a strictly positive δ can improve generalization beyond experimental conditions. In particular, for cell lines that DeepCINET encountered during training, a positive δ improves the performance of the model in most cases. More importantly DeepCINET is able to generalize to new samples better than strong baselines in a number of cases.

These results show that substantial care needs to be taken when building predictive precision oncology models from cancer cell line viability screens. We have provided evidence that explicitly accounting for phenotypic measurement noise can improve model performance, supporting the need for careful benchmarking of existing supervised models as they have been fit to data with noisy labels. Given the inconsistency within studies shown here as well as across studies in our prior work [7,8], it is imperative for models to be evaluated across multiple CCL viability screens. Within the context of precision oncology, DeepCINET presents a meaningful advance to developing biomarkers that are robust to experimental noise and that generalize beyond a single cell line screen. Currently, DeepCINET is only an advance in pre-clinical drug modeling. We plan to investigate the utility of DeepCINET to predicting real world clinical patient data in the future.

This study has several limitations. First, we have only tested DeepCINet on cancer cell lines which limits the immediate clinical utility of the model. We hope to evaluate DeepCINet on patient data in future work. Additionally, while we used gene expression as input for the prediction of drug response, DeepCINET can be applied to other types of omics measurements, such as mutation, copy number, ATAC-Seq, etc. It is possible that for some of the drugs analyzed in this paper the right type of molecular measurement for predicting genotype-phenotype relationships is not gene expression. Another limitation is that we only considered pancancer modelling. Building DeepCINET models that are cancer-type-specific may yield better results depending on the available sample size. This approach would also allow for the inclusion of cancer specific datasets such as GRAY[29] or NCI-Sarcoma[30]. Further, we only tested DeepCINET with a fixed architecture. It is possible that the model has either too high or insufficient capacity for some drugs and is well suited for others. Exploring the space of architectures could lead to improved performance. Lastly, there are cases where DeepCINET is outperformed by comparatively simpler models. As such DeepCINET does not represent a clear improvement over the state of the art in every case.

Several promising directions remain for future research. A natural next step is to evaluate DeepCINet using additional molecular features, including DNA methylation, somatic mutations, and copy-number variation. All of these data modalities are known to be implicated in drug response[31,32],[33]. Given the rise in multi-modal approaches to drug response prediction [34,35] it is possible that replacing any one omic measurement type with a multi-modal embedding from one of many recently developed foundation models [36][37][36] could lead to greater improvements in DeepCINET. We also hope to more precisely characterize when δ is helpful and how to select it for optimal generalization. The question of how to properly select δ in a principled way is left for future work. We could also consider the related problem of ordering drugs instead of cell lines, that is, sorting drugs most likely to be effective against a given cell line. This would be an application of DeepCINET to the drug recommendation literature. There are existing approaches that have already been developed to push more effective drugs to the top of the list [38] [39]. Given that our results show that DeepCINET seems to provide the best performance improvement in ranking metrics, it is possible that the model as described here can be effectively applied to this domain without much modification.

In conclusion, our findings show that explicitly accounting for phenotypic assay noise can improve the robustness of drug-response prediction. DeepCINet provides a promising framework for reducing the influence of unreliable response measurements and supporting more reliable multivariable biomarker discovery. Future work should evaluate its generalizability across additional datasets and molecular data types.

## Materials and Methods

### Data Collection & Preprocessing

Data for the Cancer Therapeutics Response Portal (CTRPv2), Cancer Cell Line Encyclopedia (CCLE), Genomics of Drug Sensitivity in Cancer (GDSC1 and GDSC2), and the Genentech Cell Line Screening Initiative (gCSI) were downloaded from the ORCESTA database [40] and pre-processed using the PharmacoGx R package[41]. For the AAC values we used the provided ‘aac_recomputed’ field available in the downloaded objects, which are computed by fitting 3-parameter Hill curves to the original study data [41]. No truncation of dose ranges nor harmonization of AAC values between studies was performed.

Drugs were selected for analysis by subjecting the 19 compounds tested in all four viability datasets - CTRPv2, GDSC1, GDSC2, and gCSI - to two quality control filters. First, drugs which had fewer than 500 cell lines for training were excluded from further analysis. Drugs that had a correlation below 0.25 in AAC measurements in common cell lines between the training screen (CTRPv2) and any of the testing screens (gCSI, GDSC2, GDSC1) were also excluded from analysis. Drugs were described as ‘broad effect’ or ‘narrow effect’ using the methods described in our prior work.[7]

All models used gene expression measurements (RNA-seq) as input features. Input genes were selected to be those plausibly relevant to cancer biology. Specifically, a set of candidate genes was constructed by taking the union of genes in the KEGG “Pathways in Cancer” gene set[42], genes in the COSMIC Gene Census[43], as well as genes thought to be targets to the 14 analyzed drugs. The potential targets were downloaded from the Therapeutic Target Database (TTD) [44]. The final features were those that were in the candidate set and also had expression measurements in every sample in each dataset. The total dimensionality of the input was 923 genes. Expression measurements were quantified in units of log-transformed transcripts per million (TPM).[45] Concretely, if *x_ij_* is the measurement in terms of TPM then the gene expression values passed to the model were log_2_ (*xij* + 0. 001).

Area above the dose response curve (AAC) measurements for each drug and cell line combination were computed as the mean across replicates in each dataset. Cell lines were restricted to only those derived from solid tumours.

### DeepCINET Architecture

The architecture used for DeepCINET was a two layer neural network with each layer having a width of 256 neurons, Rectified Linear Unit (ReLU) activations, and single dimensional output layer that served as the cell line score. The model was trained with drop out[46] with a probability of 0.5 and batch normalization [47]. DeepCINET was built using the Pytorch library [48].

### Established Machine Learning Models

All of the baseline ML models, including kernelized ridge regression[19], elastic net[20], and random forests [21] - were built using the scikit-learn python [49]. The single variable predictors were computed by selecting the gene which had the highest mutual information[50] between its expression level and measured AAC values. The final predictor was a linear regression model that used that gene’s expression value as the only covariate. This model was also built using scikit-learn.

### Model Training

The models built with scikit-learn were fitted using the built in solvers. The hyperparameters for each of the ML models, including DeepCINET, were selected using 10-fold cross validation on the training set (Supplementary Table 4).

Every DeepCINET configuration was trained for 30 epochs using the Adam [51] optimization procedure. In all cases the learning rate was set to 0.00001. Following RankNet[9] the binary cross entropy loss was used to ensure that DeepCINET assigned higher scores to cell lines more sensitive to a given drug. In the experiments where delta was tuned we again used 10 fold cross validation.

### Evaluation Metrics

Model performance was evaluated using the well known Pearson correlation coefficient [52] and the normalized discounted cumulative gain (NDCG) [9,53]. As the latter metric is less well known, we define it below.

Let *y*_1_, *y*_2_, ···, *y_n_* be the true AAC values for *n* cell lines and, for any permutation π on the numbers 1 ··· *n*, let *y*_π(1)_, *y*_π(2)_, ···, *y*_π(*n*)_ be the AAC values sorted according to π. Further let ρ be the permutation that sorts the *y* values in decreasing order. The discounted cumulative gain at K (DCG@K) for the ordering π is computed as a weighted sum:

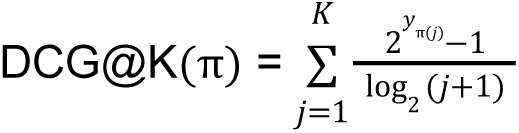

To compare between permutations the DCG@K is *normalized* by dividing by the largest possible value for the DCG@K which the value attained by computing it for the permutation ρ. Explicitly:

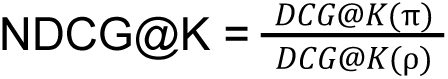

The largest possible value for NDCG is 1, which corresponds to a perfect ordering of the items by decreasing AAC. The minimum value for NDCG is 0 which corresponds to the poorest possible ordering. Machine learning models were given NDCG scores by computing the value for permutation of the AAC values given by sorting the model outputs in decreasing order. For our evaluation we set K to 5 and 10 in order to make sure we could evaluate all drugs by the same metrics. Specifically, each thanand also to keep in line with existing drug ranking literature <u>(He et al. 2020)</u>.

The robust concordance index (rCI) was computed in the manner described in [18]. Specifically for two lists of values *x* = *x*_1_, *x*_2_, ···, *x_n_* ; *y* = *y*_1_, *y*_2_, ···, *y_n_*, and a value of δ ≥ 0, we define *rCI*(*x*, *y*; δ) as the ratio of concordant pairs *CP*(*x*, *y*; δ) to valid pairs *VP*(*x*, *y*; δ).

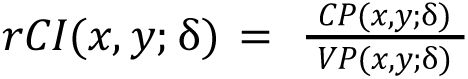

where

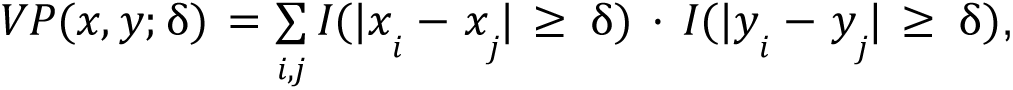

counts the number of pairs that are separated by at least δ in both lists, and

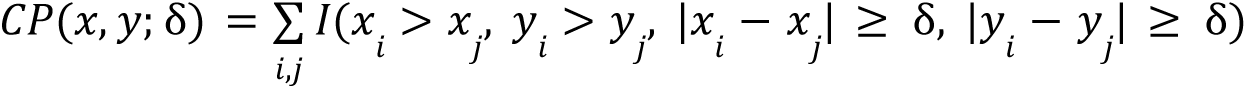

counts the number of pairs that are separated by at least δ that are ordered in the same in both *x* and *y*. Here *I* denotes the indicator function that takes the value 1 if all of its arguments are true and 0 otherwise.

## Acknowledgements

This research was supported by the NSERC: Discovery Grant RGPIN-2021-02680 and the Advanced Research Projects Agency for Health (ARPA-H), contract number 140D042590013. The views and conclusions contained in this article are those of the authors and should not be interpreted as representing the official policies, either expressed or implied, of the United States Government.

We are grateful to Kaitlyn Kobayashi and Sisira Kadambat Nair for their helpful feedback in preparing this manuscript.

## Author Contributions

JB, KLS, FK, MD, & BHK conceptualized the experiments. JB wrote the code to perform the experiments, performed the analysis, and wrote the first draft of the manuscript. KLS, BHK, and JB designed the figures. All authors reviewed the manuscript.

## Data Availability Statement

The data used in this paper are publicly available on the <u>ORCESTRA.ca</u> database and can also be accessed through the PharmacoGx R package. Using PharmacoGx we accessed the following datasets: CTRPv2 (<u>10.5281/zenodo.7826870</u>), CCLE (<u>10.5281/zenodo.3905461</u>), GDSC1 (<u>10.5281/zenodo.7829915</u>), GDSC2 (<u>10.5281/zenodo.5787145</u>), gCSI (<u>10.5281/zenodo.7829857</u>).

## Code Availability Statement

The code needed to reproduce this study can be found in the following github repository: https://github.com/bhklab/DeepConcordanceModels

## Additional Information

The authors declare that they have no competing interests.

## Supplemental Material

**Supplementary Figure 1:**
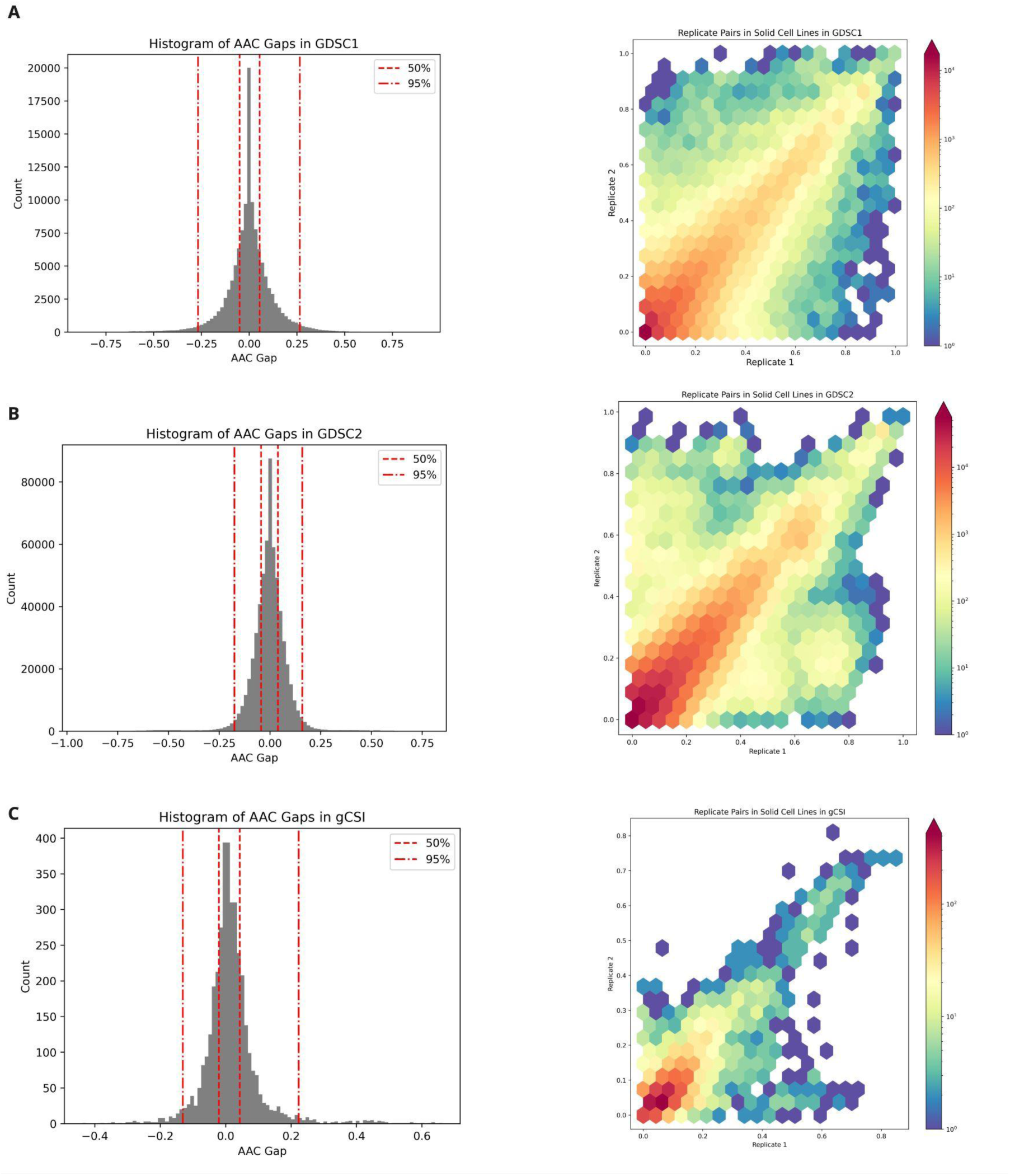
The distribution of AAC differences between replicates in gCSI and GDSC2. We show the distribution of differences in AAC measurements (top) and absolute differences (bottom) in gCSI (left) and GDSC2 (right).

**Supplementary Figure 2.**
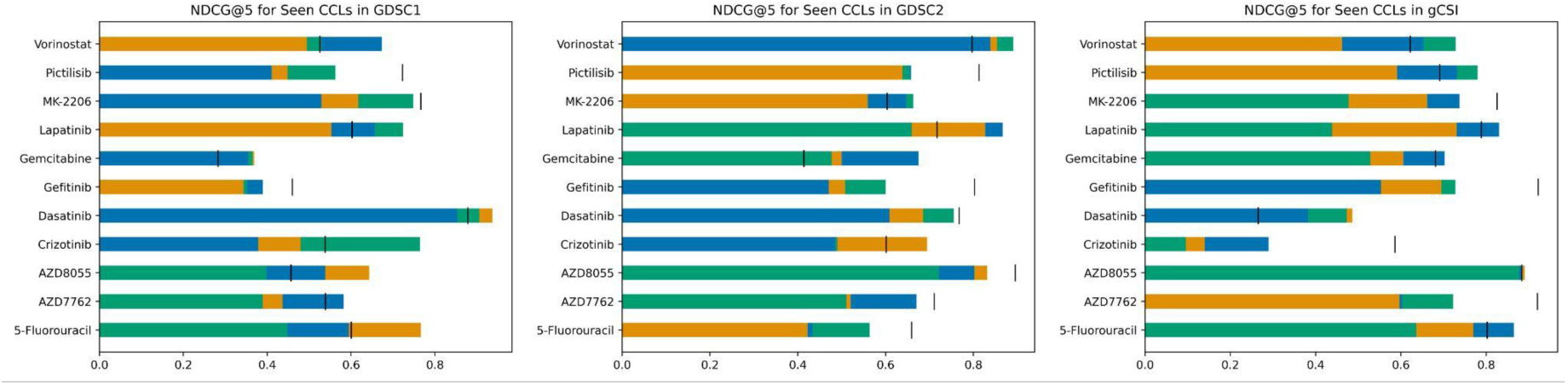
Performance as measured by NDCG@5 in seen cell lines.

**Supplementary Table 1.**
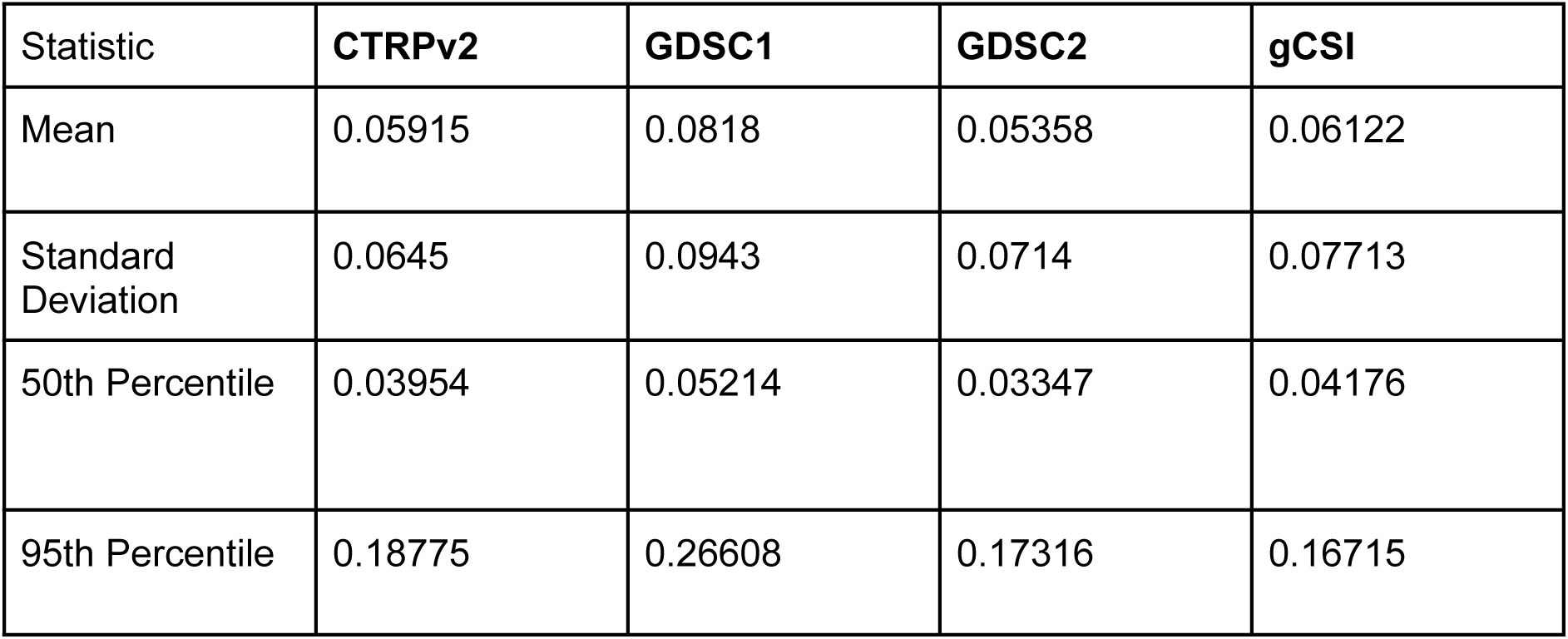
Summary Statistics of AAC Difference Distributions Within Each Dataset. We show the mean, standard deviation, 50th percentile, and 95th percentile for the distribution of absolute differences in pairs of replicates (|AAC(replicate 1)-AAC(replicate 2)|) for each of the four data sets considered.

| Statistic | **CTRPv2** | **GDSC1** | **GDSC2** | **gCSI** |
|---|---|---|---|---|
| Mean | 0.05915 | 0.0818 | 0.05358 | 0.06122 |
| Standard Deviation | 0.0645 | 0.0943 | 0.0714 | 0.07713 |
| 50th Percentile | 0.03954 | 0.05214 | 0.03347 | 0.04176 |
| 95th Percentile | 0.18775 | 0.26608 | 0.17316 | 0.16715 |

**Supplementary Table 2:** Summary Statistics of AAC Difference Distributions Within Each Dataset. We show the mean, standard deviation, 50th percentile, and 95th percentile for the distribution of differences in pairs of replicates (AAC(replicate 1)-AAC(replicate 2)) for each of the four datasets considered.

| **Statistic** | **CTRPv2** | **GDSC1** | **GDSC2** | **gCSI** |
|---|---|---|---|---|
| Mean | -0.00866 | 0.00076 | 0.01671 | -0.00394 |
| Standard Deviation | 0.08709 | 0.12483 | 0.08769 | 0.0984 |
| 50th Percentile | -0.00433 | 0.0 | 0.00839 | 0.0 |
| 95th Percentile | 0.13113 | 0.19811 | 0.1475 | 0.12016 |

**Supplementary Table 2:** The value of δ selected for each drug by 10-fold cross validation.

| **Drug** | Value for δ Selected by Cross Validation |
|---|---|
| 5-Fluorouracil | 0.13 |
| AZD7762 | 0.0 |
| AZD8055 | 0.05 |
| Crizotinib | 0.04 |
| Dasatinib | 0.11 |
| Gefitinib | 0.17 |
| Gemcitabine | 0.15 |
| Lapatinib | 0.2 |
| MK-2206 | 0.13 |
| Pictilisib | 0.17 |
| Vorinostat | 0.12 |

**Supplementary Table 3:**
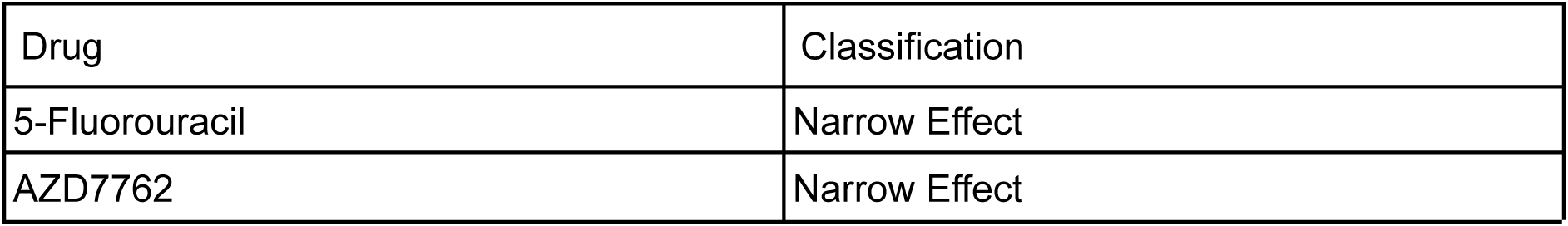
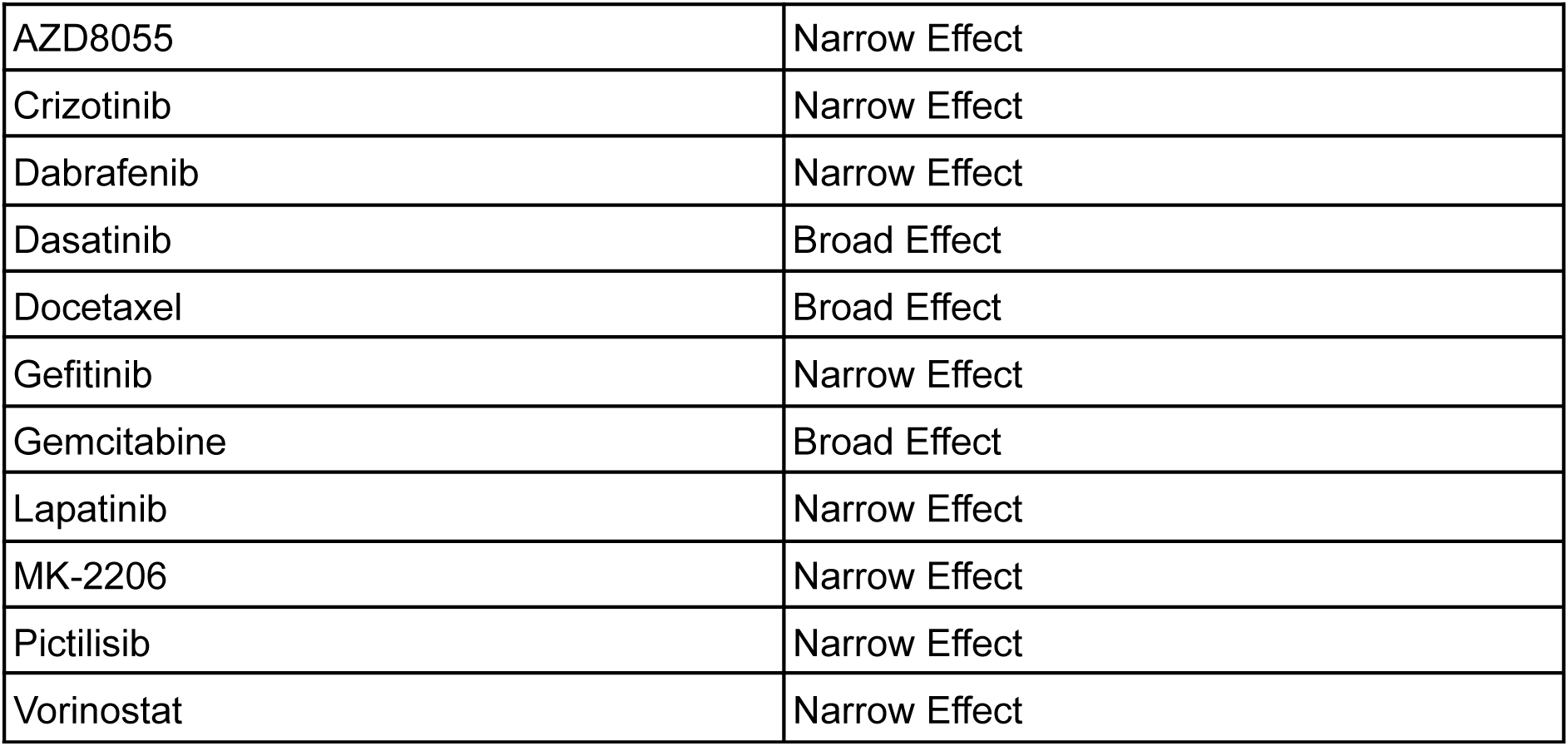
Classification of Drugs Range of Effect Using the AAC Distributions.

**Supplementary Table 4.** Hyperparameter Ranges Searched during Cross Validation.

| **Model** | **Hyperparameter** | **Range** |
|---|---|---|
| Elastic Net | Alpha | 50 values in a logarithmic grid between 0.01 and 100 |
|  | L1 Ratio | {0.1, 0.2, 0.5, 0.8, 0.85, 0.9, 0.95, 0.99, 1} |
| Kernelized Ridge Regression | Alpha | 100 equally spaced values between 0.01 and 10. |
| Random Forest | Max Features | {“sqrt”,”log2”, 1.0} |
|  | Num Estimators | {1,5,10,25,50,75,100} |
| DeepCINET | Delta (δ) | 0,0.01, …, 0.2 |
|  | Weight Decay | [0,0.01,0.1,1,5] |

**Supplementary Figure 3.**
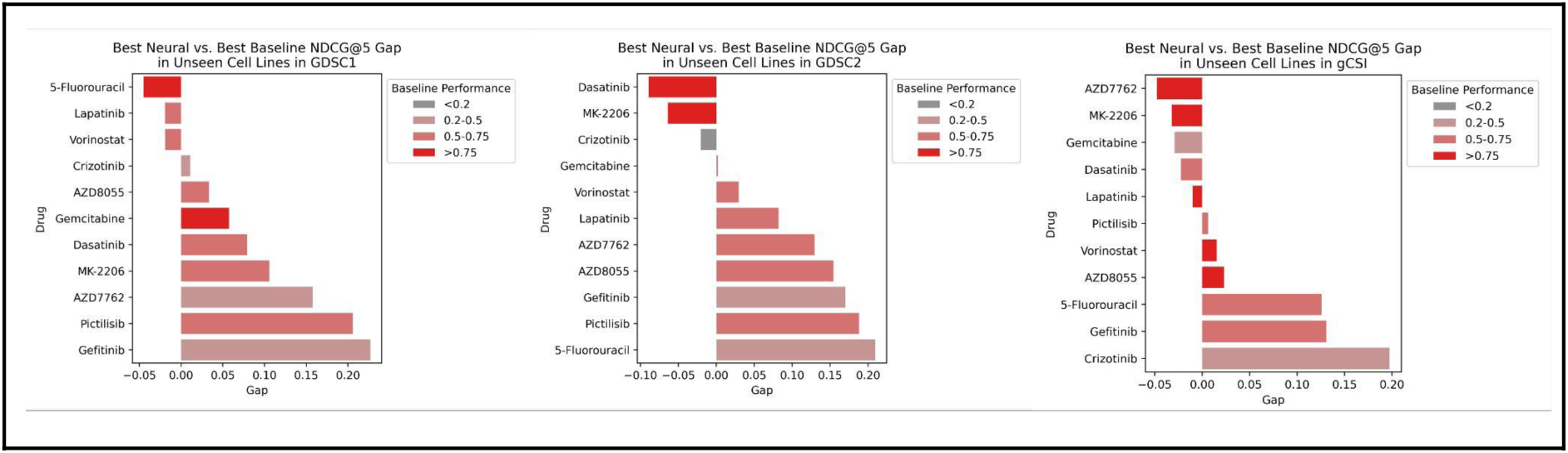
Performance as Measured by NDCG@5 in Seen Cell Lines.

**Supplementary Table 5:** The Number of Valid Pairs Used to Train Each DeepCINET Configuration.

| **Drug** | **0** | **50%** | **95%** | **CV** |
|---|---|---|---|---|
| 5-Fluorouracil | 198,765 | 138,775 | 20,341 | 46,523 |
| AZD7762 | 207,046 | 162,602 | 43,794 | 207,046 |
| AZD8055 | 205,120 | 172,366 | 69,833 | 163,982 |
| Crizotinib | 192,510 | 130,592 | 16,229 | 129,946 |
| Dasatinib | 202,566 | 176,121 | 92,551 | 133,074 |
| Gefitinib | 179,700 | 137,268 | 38,509 | 44,703 |
| Gemcitabine | 153,735 | 137,142 | 80,294 | 93,573 |
| Lapatinib | 171,991 | 125,068 | 38,773 | 35,084 |
| MK-2206 | 170,236 | 119,091 | 23,431 | 46,776 |
| Pictilisib | 202,566 | 158,123 | 40,369 | 49,130 |
| Vorinostat | 189,420 | 128,084 | 12,095 | 41,235 |

**Supplementary Table 6:** The Number of Cell Lines, AAC Range, and the tested Dose Range in CTRPv2 for Each Drug Considered.

| **CTRPv2** |  |  |  |
| **Drug** | **Num. Cell Lines** | **AAC Range** | **Tested Dose Range (µM)** |
|---|---|---|---|
| 5-Fluorouracil | 631 | [0.0,0.48] | 0.0041, 0.0081, 0.016, 0.032, 0.065, 0.13, 0.26, 0.52, 1, 2.1, 4.2, 8.3, 17, 33, 66, 130 |
| AZD7762 | 644 | [0.03,0.805] | 0.002, 0.0041, 0.0081, 0.016, 0.032, 0.065, 0.13, 0.26, 0.52, 1, 2.1, 4.2, 8.3, 17, 33, 66 |
| AZD8055 | 641 | [0.0,0.76] | 0.002, 0.0041, 0.0081, 0.016, 0.032, 0.065, 0.13, 0.26, 0.52, 1, 2.1, 4.2, 8.3 ,17, 33, 66 |
| Crizotinib | 621 | [0.0,0.728] | 0.002, 0.0081, 0.016, 0.032, 0.065, 0.13, 0.26, 0.52, 1, 2.1, 4.2, 8.3, 17, 33, 66 |
| Dasatinib | 637 | [0.0,0.923] | 0.002, 0.0041, 0.0081, 0.016, 0.032, 0.065, 0.13, 0.26, 0.52, 1, 2.1, 4.2, 8.3, 17, 33, 66 |
| Gefitinib | 600 | [0.0,0.849] | 0.002, 0.0041, 0.0081, 0.016, 0.032, 0.065, 0.13, 0.26, 0.52, 1, 2.1, 4.2, 8.3, 17, 33 |
| Gemcitabine | 555 | [0.0,0.868] | 0.001, 0.002, 0.0041, 0.0081, 0.016, 0.032, 0.065, 0.13, 0.26, 0.52, 1, 2.1, 4.2, 8.3, 17, 33 |
| Lapatinib | 587 | [0.0,0.644] | 0.002, 0.0041, 0.0081, 0.016, 0.032, 0.065, 0.13, 0.26, 0.52, 1, 2.1, 4.2, 8.3, 17, 33, 66 |
| MK-2206 | 584 | [0.0,0.543] | 0.001, 0.002, 0.0041, 0.0081, 0.016, 0.032, 0.065, 0.13, 0.26, 0.52, 1, 2.1, 4.2, 8.3, 17, 33 |
| Pictilisib | 637 | [0.0,0.886] | 0.002, 0.0041, 0.0081, 0.016, 0.032, 0.065, 0.13, 0.26, 0.52, 1, 2.1, 4.2, 8.3, 17, 33, 66 |
| Vorinostat | 616 | [0.0,0.62] | 0.001, 0.002, 0.0041, 0.0081, 0.016, 0.032, 0.065, 0.13, 0.26, 0.52, 1, 2.1, 4.2, 8.3, 17, 33 |

**Supplementary Table 7:** The Number of Cell Lines, AAC Range, and the tested Dose Range in GDSC1 for Each Drug Considered.

| GDSC1 |  |  |  |
| **Drug** | **Num. Cell Lines** | **AAC Range** | **Tested Dose Range (µM)** |
|---|---|---|---|
| 5-Fluorouracil | 325 | [0.0,0.615] | 0.125,0.25,0.5,1,2,4, 8,16,32 |
| AZD7762 | 332 | [0.0,0.799] | 0.007, 0.015, 0.031, 0.0625, 0.125, 0.25, 0.5, 1, 2 |
| AZD8055 | 326 | [0.0,0.599] | 0.007, 0.015, 0.031, 0.0625, 0.125, 0.25, 0.5, 1, 2 |
| Crizotinib | 137 | [0.0,0.404] | 0.0078125, 0.015625, 0.03125, 0.0625, 0.125, 0.25, 0.5, 1, 2 |
| Dasatinib | 137 | [0.0,0.894] | 0.005, 0.01, 0.02, 0.04, 0.08, 0.16, 0.32, 0.64, 1.28 |
| Gefitinib | 333 | [0.0,0.473] | 0.00195, 0.00391, 0.00781, 0.015625, 0.03125, 0.0625, 0.125, 0.25, 0.5 |
| Gemcitabine | 295 | [0.0,0.974] | 0.004, 0.008, 0.016, 0.032, 0.064, 0.128, 0.256, 0.512, 1.024 |
| Lapatinib | 136 | [0.0,0.584] | 0.0078125, 0.015625, 0.03125, 0.0625, 0.125, 0.25, 0.5, 1, 2 |
| MK-2206 | 298 | [0.0,0.625] | 0.015625, 0.03125, 0.0625, 0.125, 0.25, 0.5, 1, 2, 4 |
| Pictilisib | 335 | [0.0,0.553] | 0.015625, 0.03125, 0.0625, 0.125, 0.25, 0.5, 1, 2, 4 |
| Vorinostat | 334 | [0.0,0.692] | 0.0390625, 0.078125, 0.15625, 0.3125, 0.625, 1.25, 2.5, 5,10 |

**Supplementary Table 8:** The Number of Cell Lines, AAC Range, and the tested Dose Range in GDSC2 for Each Drug Considered.

| GDSC2 |  |  |  |
| Drug | Num. Cell Lines | AAC Range | Tested Dose Range (µM) |
|---|---|---|---|
| 5-Fluorouracil | 260 | [0.0,0.328] | 0.0041, 0.0081, 0.016, 0.032, 0.065, 0.13, 0.26, 0.52, 1, 2.1, 4.2, 8.3, 17, 33, 66, 130 |
| AZD7762 | 243 | [0.028,0.695] | 0.002, 0.0041, 0.0081, 0.016, 0.032, 0.065, 0.13, 0.26, 0.52, 1, 2.1, 4.2, 8.3, 17, 33, 66 |
| AZD8055 | 40 | [0.028,0.625] | 0.002, 0.0041, 0.0081, 0.016, 0.032, 0.065, 0.13, 0.26, 0.52, 1, 2.1, 4.2, 8.3, 17, 33, 66 |
| Crizotinib | 234 | [0.0,0.331] | 0.002, 0.0041, 0.0081, 0.016, 0.032, 0.065, 0.13, 0.26, 0.52, 1, 2.1, 4.2, 8.3, 17, 33, 66 |
| Dasatinib | 242 | [0.0,0.538] | 0.002, 0.0041, 0.0081, 0.016, 0.032, 0.065, 0.13, 0.26, 0.52, 1, 2.1, 4.2, 8.3, 17, 33, 66 |
| Gefitinib | 239 | [0.0,0.387] | 0.002,0.0041, 0.0081, 0.016, 0.032, 0.065, 0.13, 0.26, 0.52, 1, 2.1, 4.2, 8.3, 17, 33, 66 |
| Gemcitabine | 243 | [0.003,0.889] | 0.001, 0.002, 0.0081, 0.016, 0.032, 0.065, 0.13, 0.26, 0.52, 1, 2.1, 4.2, 8.3, 17, 33 |
| Lapatinib | 240 | [0.0,0.624] | 0.002, 0.0041, 0.0081, 0.016, 0.032, 0.065, 0.26, 0.52, 1, 2.1, 4.2, 8.3, 17, 33 |
| MK-2206 | 245 | [0.0,0.486] | 0.001, 0.002, 0.0041, 0.0081, 0.016, 0.032, 0.065, 0.13, 0.26, 0.52, 1, 2.1, 4.2, 8.3, 17, 33 |
| Pictilisib | 243 | [0.0,0.463] | 0.002, 0.0041, 0.0081, 0.016, 0.032, 0.065, 0.13, 0.26, 0.52, 1, 2.1, 4.2, 8.3, 17, 33, 66 |
| Vorinostat | 240 | [0.041,0.514] | 0.001, 0.002, 0.0041, 0.0081, 0.016, 0.032, 0.065, 0.13, 0.26, 0.52, 1, 2.1, 4.2, 8.3,, 17, 33 |

**Supplementary Table 9:** The Number of Cell Lines, AAC Range, and the tested Dose Range in GDSC2 for Each Drug Considered.

| **gCSI** |  |  |  |
| **Drug** | **Num. Cell Lines** | **AAC Range** | **Tested Dose Range (µM)** |
|---|---|---|---|
| 5-Fluorouracil | 92 | [0.081,0.647] | 0.03, 0.09, 0.27, 0.82, 2.47,7.41, 22.22, 66.67, 200 |
| AZD7762 | 71 | [0.171,0.753] | 0.0030, 0.0091, 0.0274, 0.0823, 0.2469, 0.7407, 2.2222, 6.6667, 20 |
| AZD8055 | 71 | [0.279,0.57] | 0.0030, 0.0091, 0.0274, 0.0823, 0.2469, 0.7407, 2.2222, 6.6667, 20 |
| Crizotinib | 344 | [0.0,0.424] | 0.00015, 0.00045, 0.0014, 0.0041, 0.0124, 0.0371, 0.1111, 0.3333, 1 |
| Dasatinib | 137 | [0.0,0.243] | 1.5-05, 4.5E-05, 0.0001, 0.0004, 0.0012, 0.0037, 0.0111, 0.0333, 0.1 |
| Gefitinib | 71 | [0.0,0.459] | 0.0003, 0.0009, 0.0027, 0.0082, 0.0246, 0.0740, 0.2222, 0.6666, 2 |
| Gemcitabine | 230 | [0.0,0.807] | 0.0001, 0.0004, 0.0014, 0.0041, 0.0123, 0.037, 0.1111, 0.3333, 1 |
| Lapatinib | 341 | [0.0,0.49] | 0.0030, 0.0091, 0.0274, 0.0823, 0.2469, 0.7407, 2.2222, 6.6666, 20 |
| MK-2206 | 71 | [0.001,0.265] | 0.0001, 0.0004, 0.0013, 0.0015, 0.0041, 0.0045, 0.0123, 0.0137, 0.0370, 0.0411, 0.1111, 0.1235, 0.3333, 0.3703, 1, 1.1111, 3.3333, 10 |
| Pictilisib | 354 | [0.0,0.462] | 0.0015, 0.0045, 0.0137, 0.0412, 0.1237, 0.3712, 1.1111, 3.3333, 10 |
| Vorinostat | 344 | [0.023,0.458] | 0.0015, 0.0046, 0.0137, 0.0412, 0.1237, 0.3712, 1.1111, 3.3333, 10 |

**Supplementary Figure 4:**
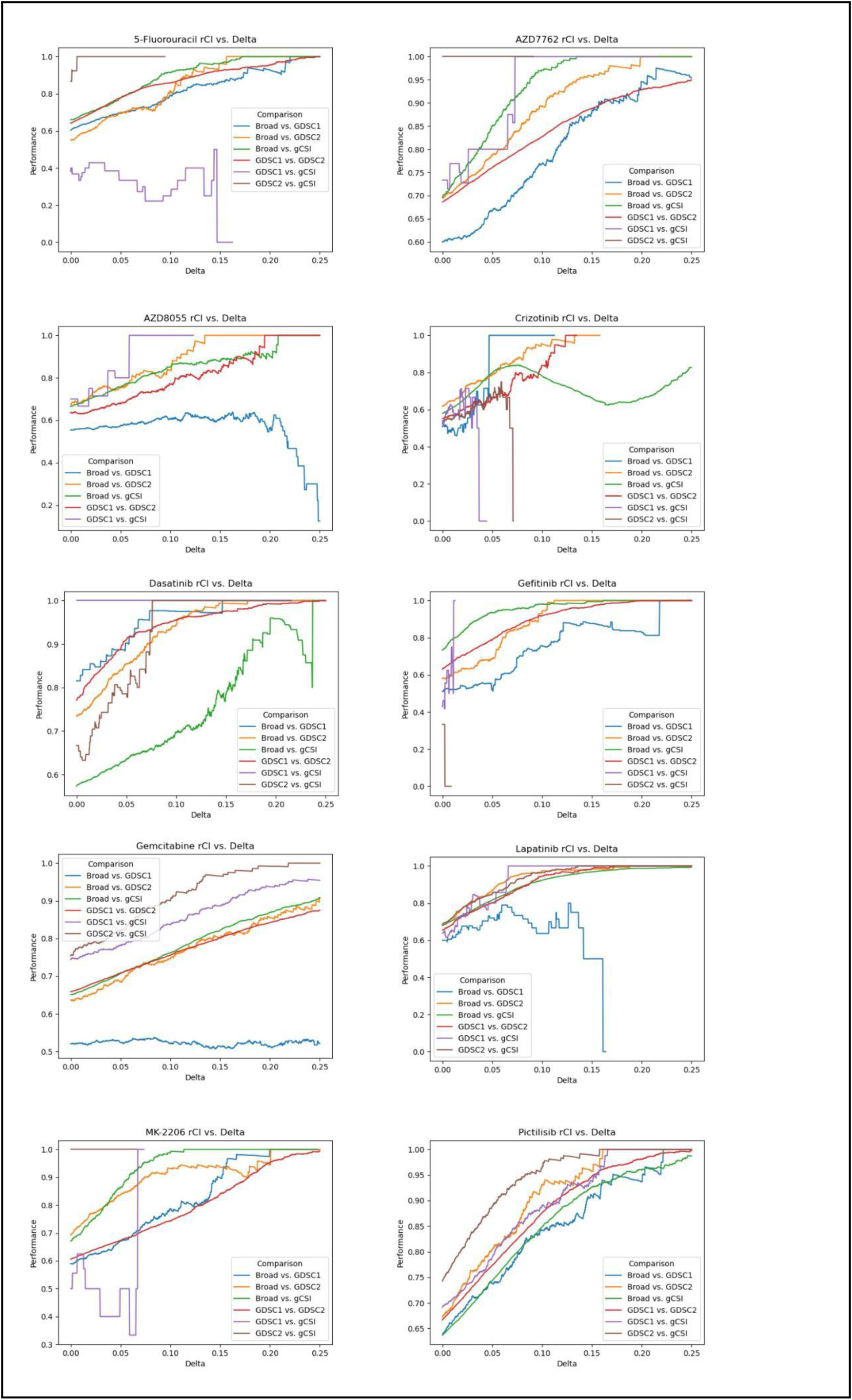
Relationship between the robust concordance index (rCI) and delta in 10 of the 11 drugs considered. Vorinostat is present in the main manuscript.

